# Deep coverage whole genome sequences and plasma lipoprotein(a) in individuals of European and African ancestries

**DOI:** 10.1101/225169

**Authors:** Seyedeh M. Zekavat, Sanni Ruotsalainen, Robert E. Handsaker, Maris Alver, Jonathan Bloom, Tim Poterba, Cotton Seed, Jason Ernst, Mark Chaffin, Jesse Engreitz, Adolfo Correa, Andres Metspalu, Veikko Salomaa, Manolis Kellis, Mark J. Daly, James G. Wilson, Benjamin M. Neale, Steven McCaroll, Ida Surakka, Tonu Esko, Andrea Ganna, Samuli Ripatti, Sekar Kathiresan, Pradeep Natarajan, NHLBI TOPMed Lipids Working Group

**Author notes:** Contributed equally (Tonu Esko, Andrea Ganna, Samuli Ripatti, Sekar Kathiresan, Pradeep Natarajan).

## Abstract

Lipoprotein(a), Lp(a), is a modified low-density lipoprotein particle where apolipoprotein(a) (protein product of the *LPA* gene) is covalently attached to apolipoprotein B. Lp(a) is a highly heritable, causal risk factor for cardiovascular diseases and varies in concentrations across ancestries. To comprehensively delineate the inherited basis for plasma Lp(a), we performed deep-coverage whole genome sequencing in 8,392 individuals of European and African American ancestries. Through whole genome variant discovery and direct genotyping of all structural variants overlapping *LPA*, we quantified the 5.5kb kringle IV-2 copy number (KIV2-CN), a known *LPA* structural polymorphism, and developed a model for its imputation. Through common variant analysis, we discovered a novel locus (*SORT1*) associated with Lp(a)-cholesterol, and also genetic modifiers of KIV2-CN. Furthermore, in contrast to previous GWAS studies, we explain most of the heritability of Lp(a), observing Lp(a) to be 85% heritable among African Americans and 75% among Europeans, yet with notable inter-ethnic heterogeneity. Through analyses of aggregates of rare coding and non-coding variants with Lp(a)-cholesterol, we found the only genome-wide significant signal to be at a non-coding *SLC22A3* intronic window also previously described to be associated with Lp(a); however, this association was mitigated by adjustment with KIV2-CN. Finally, using an additional imputation dataset (N=27,344), we performed Mendelian randomization of *LPA* variant classes, finding that genetically regulated Lp(a) is more strongly associated with incident cardiovascular diseases than directly measured Lp(a), and is significantly associated with measures of subclinical atherosclerosis in African Americans.

## MAIN TEXT

Lipoprotein(a), Lp(a), is a circulating lipoprotein comprised of a modified low-density lipoprotein (LDL) particle covalently bonded to apolipoprotein(a), apo(a)^1–3^. The apo(a) protein contains an inactive protease domain, kringle V domain, and ten kringle IV domains, including an extremely polymorphic kringle IV 2 copy number (KIV2-CN)^3^, a large region spanning 5.5kb which consists of a pair of exons repeating between 5 to over 40 times per chromosome^4^. Increased KIV2-CN results in increased apo(a) size which is inversely associated with plasma Lp(a) levels due to altered protein folding, transport, and secretion^5^. Twin studies have suggested that Lp(a) is highly heritable, with up to 90% heritability in both African and European populations^6–10^. However, the most recent genome-wide association studies have only explained approximately half of the genetic heritability^11^. Epidemiologic studies and genetic analyses in European and Asian populations have causally linked Lp(a) concentrations with atherosclerotic cardiovascular disease, independent of other plasma lipids including LDL cholesterol^12–15^. As a result, Lp(a) has emerged as a promising therapeutic target for atherosclerotic cardiovascular diseases.

Plasma Lp(a) distributions vary significantly among ethnicities but these differences are not explained by known differential KIV2-CN distributions between the ethnicities and are posited to be related to primary sequence^16^. Additionally, studies suggest that apo(a) isoform and Lp(a) concentration may have differential effects on coronary heart disease (CHD) odds^14^; however, distinguishing isoform-independent genetic effects on Lp(a) has required separate genotyping strategies, typically qPCR^17^, in addition to genotyping single nucleotide polymorphisms (SNPs). Deep-coverage (>20X) whole genome sequencing (WGS) provides the opportunity to determine the full range of genomic variation that influences Lp(a) concentration and isoform size, across the allele frequency spectrum and variant type among diverse individuals.

Here, we used deep-coverage WGS in 2,284 Estonians, 2,690 Finnish individuals, and 3,418 African Americans to ascertain SNPs and indels across the genome, and structural variants at *LPA*, including KIV2-CN. We performed: 1) structural variant association analyses; 2) common variant association; 3) rare variant association in coding and non-coding sequence; and 4) Mendelian randomization analyses. Our goals were three-fold: 1) to understand the full spectrum of genetic variation influencing Lp(a) and Lp(a)-cholesterol (Lp(a)-C); 2) to compare genetic differences between Europeans and African Americans; and 3) to determine the phenotypic consequences of *LPA* variant classes on incident clinical events and subclinical measures (**Fig. 1**).

**Fig 1.**
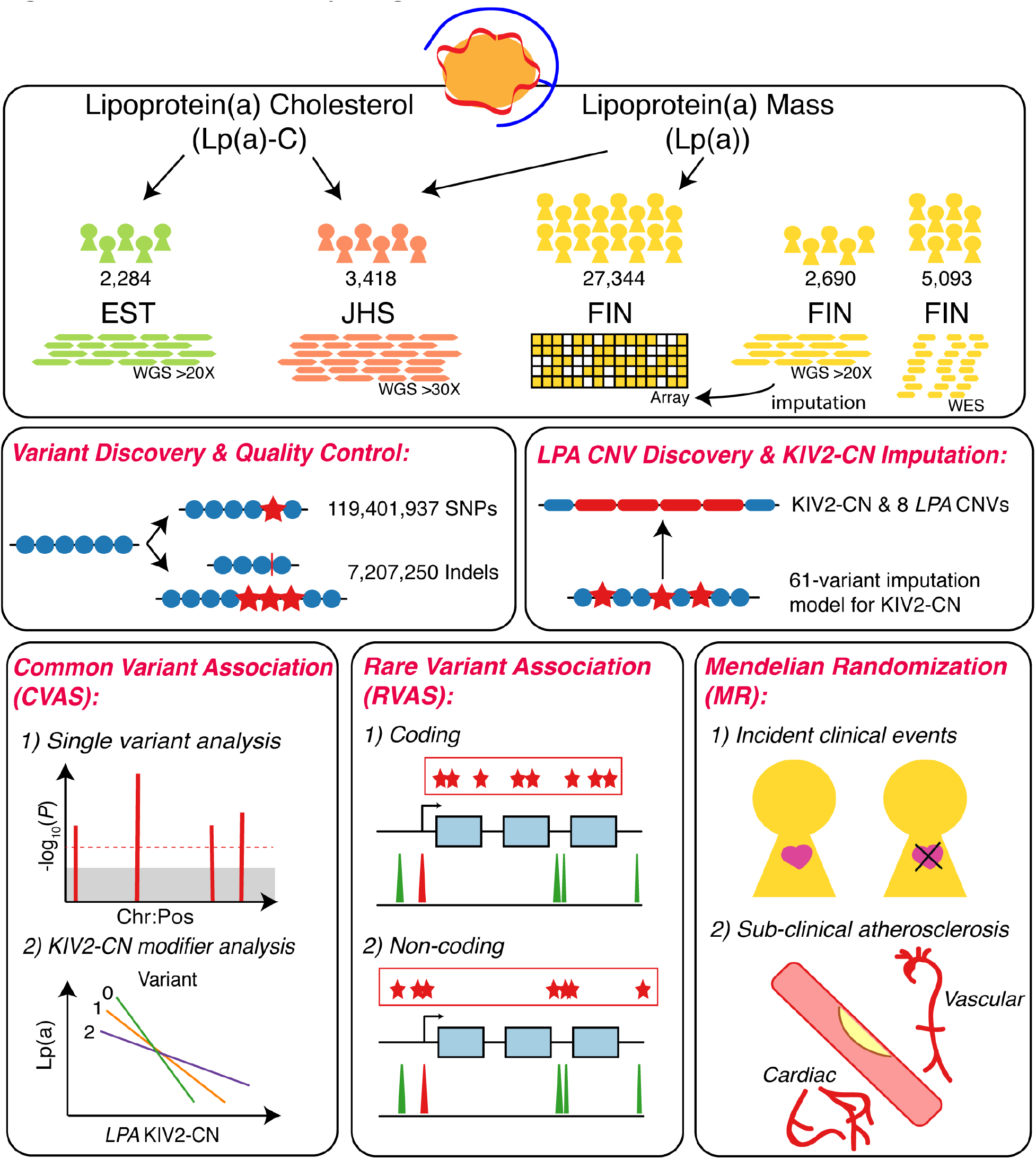
Schema of overall study design. Analyses were stratified by phenotype, Lp(a) (mass) and Lp(a)-C, where available. Lp(a)-C analyses were performed using the following individuals with WGS data: 2,284 individuals from the Estonian Biobank (EST) and 3,418 individuals from Jackson Heart Study (JHS). Lp(a) mass analyses were performed using the same Jackson Heart Study participants as well as array-derived genotypes from 27,344 Finnish FINRISK (FIN) individuals with imputation performed using 2,690 FIN individuals with WGS and 5,093 FIN individuals with WES. After quality control filters, 119,401,837 SNPs and 7,207,350 indels were discovered genome-wide across individuals analyzed. Structural variant discovery at the LPA locus was performed, finding KIV2-CN and 8 additional rare CNVs. An imputation model was developed to impute KIV2-CN using 60 LPA-locus variants. Three overarching analyses were subsequently performed: 1) Common variant analyses, 2) Rare variant analyses, and 3) Mendelian randomization. Among common and low-frequency variants with MAF > 0.1%, we performed single variant analysis, and separately, analyzed genetic modifiers of KIV2-CN’s effect on Lp(a) and Lp(a)-C concentrations. We also performed rare variant analyses, aggregating rare variants (MAF < 1%) in 1) coding sequence and 2) putative functional noncoding sequence, and associated with Lp(a)-C. Lastly, we performed Mendelian randomization, using different classes of variants associated with Lp(a) as genetic instruments and associating these with incident clinical cardiovascular events in FIN and prevalent subclinical atherosclerosis in JHS. CNV = copy number variant; EST = Estonian biobank; FIN = FINRISK; JHS = Jackson Heart Study; KIV2-CN = kringle IV-2 copy number; Lp(a) = lipoprotein(a); Lp(a)-C = lipoprotein(a) cholesterol; MAF = minor allele frequency; MR = Mendelian randomization

### Whole genome sequencing and baseline characteristics

A total of 8,392 participants underwent deep-coverage (mean attained 33X coverage) WGS: 3,418 African Americans from the Jackson Heart Study (JHS) as part of the NIH/NHLBI Trans-Omics for Precision Medicine (TOPMed) program, 2,284 Europeans from the Estonian Biobank (EST), and 2,690 Europeans from the Finland FINRISK study (FIN) (**Supplementary Fig. 1**). FIN WGS and whole exome sequences were used to impute into 27,344 Finnish array data for analyses. Following quality control (**Supplementary Table 1**), a total of 119.4M SNPs and 7.2M indels were discovered across EST WGS, JHS WGS, and FIN imputation datasets analyzed (**Supplementary Fig. 2–3, Supplementary Table 2**).

We obtained both Lp(a) and Lp(a)-C where available. 4,767 individuals from EST and JHS WGS with Lp(a)-C available and 9,272 individuals from the JHS WGS and FIN imputation dataset with Lp(a) available were included in analyses requiring these phenotypes. Lp(a)-C values were quantified using the Vertical Autoprofile (VAP) method, which measures cholesterol concentration via densitometry^18,19^. Lp(a) values were quantified using an immunoassay-based method sensitive to the entire mass of the Lp(a) particle. Median Lp(a) levels in JHS (median (IQR) 46 (24-79) mg/dL) were nearly ten times higher than in FIN (5 (2-10) mg/dL), while the Lp(a)-C distribution was similar between EST (7 (5-9) mg/dL) and JHS (7 (5-11) mg/dL) (**Supplementary Table 3, Supplementary Fig. 4a,b**). We note that we are particularly well powered to detect genetic differences between individuals of African and European ancestry since Finnish individuals are known to have the lowest Lp(a) concentration amongst different European populations^20^, thus explaining why we have a 10-fold difference between JHS and FIN Lp(a) concentrations as opposed to the 2–3 fold differences typically described between European and African populations^16^. Among JHS individuals with both Lp(a) and Lp(a)-C available, the concentrations between these phenotypes were highly correlated (Spearman correlation (R_s_) = 0.46, *P* = 2.4×10^−143^) (**Supplementary Fig. 5**).

### Structural variant analyses: discovery and imputation of KIV2-CN from sequence reads

Structural variants, notably KIV2-CN, at *LPA* have been previously shown to influence apo(a) size and Lp(a) concentration^17^. From the WGS data, we used GenomeSTRiP^21^ to identify and genotype 9 structural variants at the *LPA* locus (**Fig. 2a, Supplementary Table 4**), all rare except the KIV2-CN repeat. We mapped the reported 6 KIV2 repeats present in the hg19 reference genome^22^, finding that the KIV2-CN repeat occurs between positions chr6:161032565-161067901 with each repeat copy containing 5,534 – 5,546 base pairs and two coding exons (**Supplementary Fig. 6a**). The KIV2-CN (quantified as the sum of the KIV2 allelic copy number across both chromosomes) distribution is slightly different between African American (mean 38.5 (SD 7.4)) and European (mean 43.7 (SD 6.2)) ethnicities, ranging between 12.0-84.6 copies (**Supplementary Fig. 6b, Supplementary Table 5**). In earlier work, we validated Genome STRiP copy number estimates using ddPCR^23^, which establishes general accuracy for the quantified absolute copy number. To evaluate the precision of our KIV2-CN estimates, we utilized 123 pairs of siblings from JHS that were confidently identical-by-descent at both *LPA* 1Mb window haplotypes (genotype concordance > 99%), and found a robust correlation between sibling pair KIV2 copy number estimates (r^2^=0.989) (**Supplementary Fig. 7a-d**).

**Fig 2.**
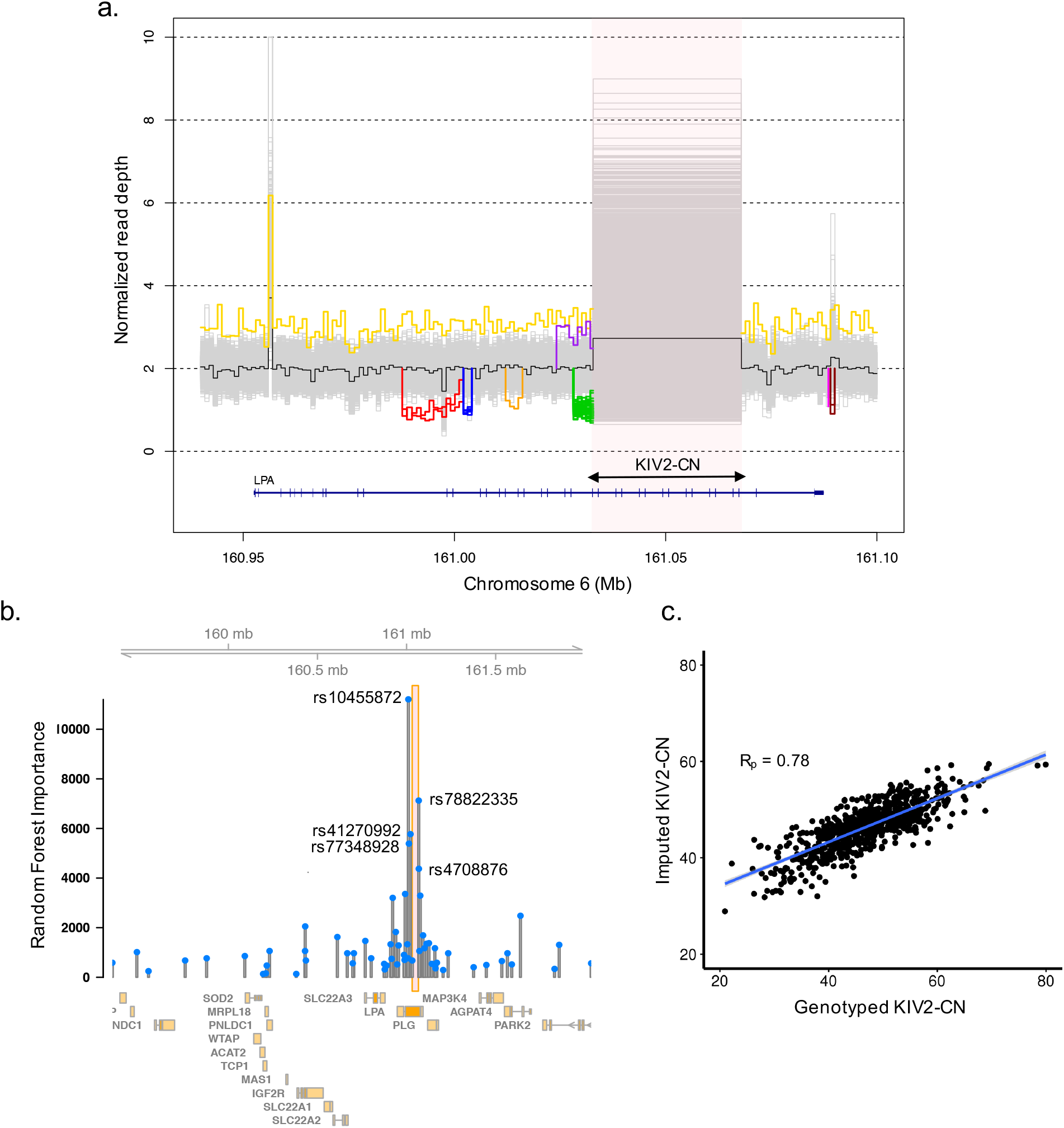
Structural variant discovery at the *LPA* locus and KIV2-CN imputation. a) Nine separate copy number variants were discovered across the EST, JHS, and FIN whole genome sequences. Here, these are shown by plotting sample-level normalized read depth against the position along the hg19 reference genome at the *LPA* locus (with the black line denoting median read depth across all individuals). The KIV2-CN is shown in the highlighted region and each unique non-gray line outside of this region depicts a discovered structural variant (described further in Supplementary Table 4). b) The random forest importance of each variant in the 61-variant KIV2-CN imputation model developed in FIN is shown against its genomic position, with KIV2-CN region highlighted and the top five rsIDs labeled. c) Correlation of directly genotyped KIV2-CN and imputed KIV2-CN from 738 FIN individuals with WGS in the validation dataset (with Pearson correlation, R_p_ = 0.78). EST = Estonian biobank; FIN = FINRISK; JHS = Jackson Heart Study; KIV2-CN = kringle IV-2 copy number

*LPA* locus variants, namely rs3798220 and rs10455872, have been previously associated with KIV2-CN^14,15^. In the FIN WGS, these two SNPs account for 12% of the variance of directly genotyped KIV2-CN. To improve KIV2-CN estimation from SNPs, we developed an imputation model using 2,215 FIN with WGS and applied it to impute KIV2-CN in the 27,344 FIN with array-derived genotypes. In the FIN WGS, we applied the least absolute shrinkage and selection operator (LASSO) across high-quality (imputation quality > 0.8) variants with minor allele frequency (MAF) > 0.1% available in the FIN imputation dataset in a 4MB window around *LPA*, which yielded a 61-variant model to impute KIV2-CN (**Supplemental Fig. 8a**). To understand the relative importance of each of these 61 variants, a random forest model was applied (**Fig. 2b, Supplementary Fig. 8b**). Our model ascribed greatest importance to rs10455872, a previously described SNP associated with KIV2-CN^14,15^. The full 61-variant model in our validation dataset explained 60% of variation in genotyped KIV2-CN (**Supplemental Table 5, Supplemental Fig. 6c, Fig. 3b**). While low frequency loss-of-function variants have been observed by us and others^24,25^ within *LPA*, removal of these carriers did not significantly alter the relationship between KIV2-CN and Lp(a) across all individuals (*P*=0.48).

**Fig 3.**
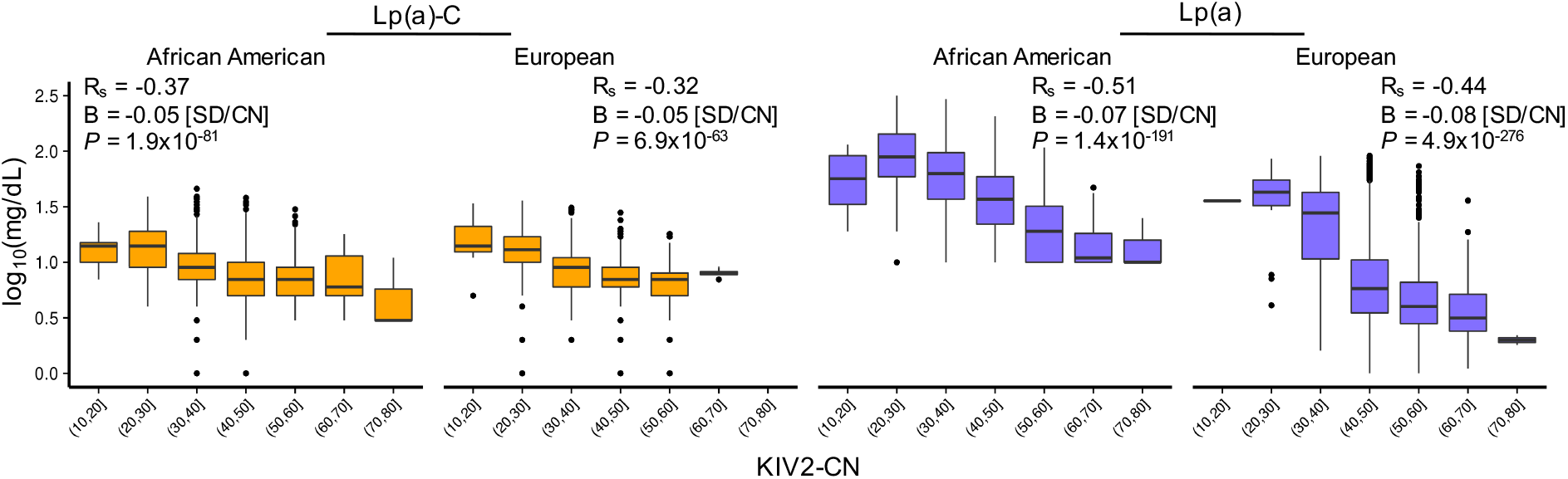
KIV2-CN association with Lp(a) phenotypes. Directly genotyped KIV2-CN (in EST and JHS) and imputed KIV2-CN (in FIN) are inversely associated with Lp(a) and Lp(a)-C. EST = Estonian biobank; FIN = FINRISK; JHS = Jackson Heart Study; KIV2-CN = kringle IV-2 copy number; Lp(a) = lipoprotein(a); Lp(a)-C = lipoprotein(a) cholesterol; R_p_ = Pearson correlation; R_S_ = Spearman correlation

We confirmed that both directly genotyped and imputed KIV2-CN were negatively associated with Lp(a)-C (-0.05 SD/CN, *P* < 1×10^−61^) and Lp(a) (−0.07 to −0.08 SD/CN, *P* < 1×10^−190^), across African American and European ethnicities (Fig. 3c). KIV2-CN alone explained 18% (Europeans) to 26% (African Americans) of variation in Lp(a), and for Lp(a)-C explained 14% of variation in both ethnicities. Introduction of 1/KIV2-CN to the multivariable model did not improve model fit for the relationship between KIV2-CN and Lp(a) (*P*=0.16).

We sought to also determine whether combinations of summed KIV2-CN alleles equivalent to the same total had the same relationship with KIV2-CN. We observed that the relationship of homozygous KIV2-CN alleles (from 59 FIN individuals 95% homozygous-by-descent at the *LPA* locus) to Lp(a) was similar to the remaining association observed across all others (*P* = 0.21).

### Common variant association: Single variant analyses

To identify additional genomic variants associated with Lp(a) and Lp(a)-C, we performed genome-wide common variant (MAF > 0.1%) association analyses using a linear mixed model, conditioning on KIV2-CN. Association was performed at the cohort-level and followed by trans ethnic meta-analysis. We analyzed a total of 32,695,476 variants for Lp(a)-C and 31,652,301 variants for Lp(a), identifying common variants at 3 loci at conventional genome-wide significance (*P* < 5×10^−8^) for Lp(a)-C at *LPA* (rs140570886, *P* = 3.3×10^−30^), *CETP* (rs247616, *P* = 6.1×10^−10^), and *SORT1* (rs12740374 *P* = 1.0×10^−21^), and 2 genome-wide significant loci for Lp(a) at *LPA* (rs6938647, *P* = 4.7×10^−129^), and *APOE* (rs7412, *P* = 1.3×10^−23^) (**Supplementary Fig. 9-11; Supplementary Tables 7, 8**).

The lead *SORT1* locus variant, rs12740374, has been previously causally associated with LDL cholesterol^26^. Here, Lp(a)-C association for rs12740374 was not substantially altered conditioned on either LDL cholesterol (**Fig. 4a**) or apolipoprotein B (**Supplementary Fig. 12**). Common variants at *CETP* are associated with HDL cholesterol^27^ and the lead *CETP* locus variant for Lp(a)-C, rs247616, is no longer significant after conditioning on HDL cholesterol (**Supplementary Fig. 13**). Lp(a)-C is strongly associated with HDL cholesterol (*B* = 0.41 SD Lp(a)-C/SD HDL, *P* =2.9×10^−191^); notably, HDL and Lp(a) particles have similar densities potentially influencing Lp(a)-C measurement accuracy^28^. Finally, rs7412 (*APOE* p. Arg176Cys), denoting the major APOE2 polymorphism, has been previously associated with LDL cholesterol^29^ and recently with Lp(a) in a meta-analysis^11^. The association of rs7412 with Lp(a) is diminished when conditioning on LDL cholesterol but remains strongly associated (before conditioning: *B* = −0.25 SD, *P* = 1×10^−23^, after conditioning: *B* = −0.18 SD, *P* = 5×10^−16^) (**Fig. 4b**).

**Fig 4.**
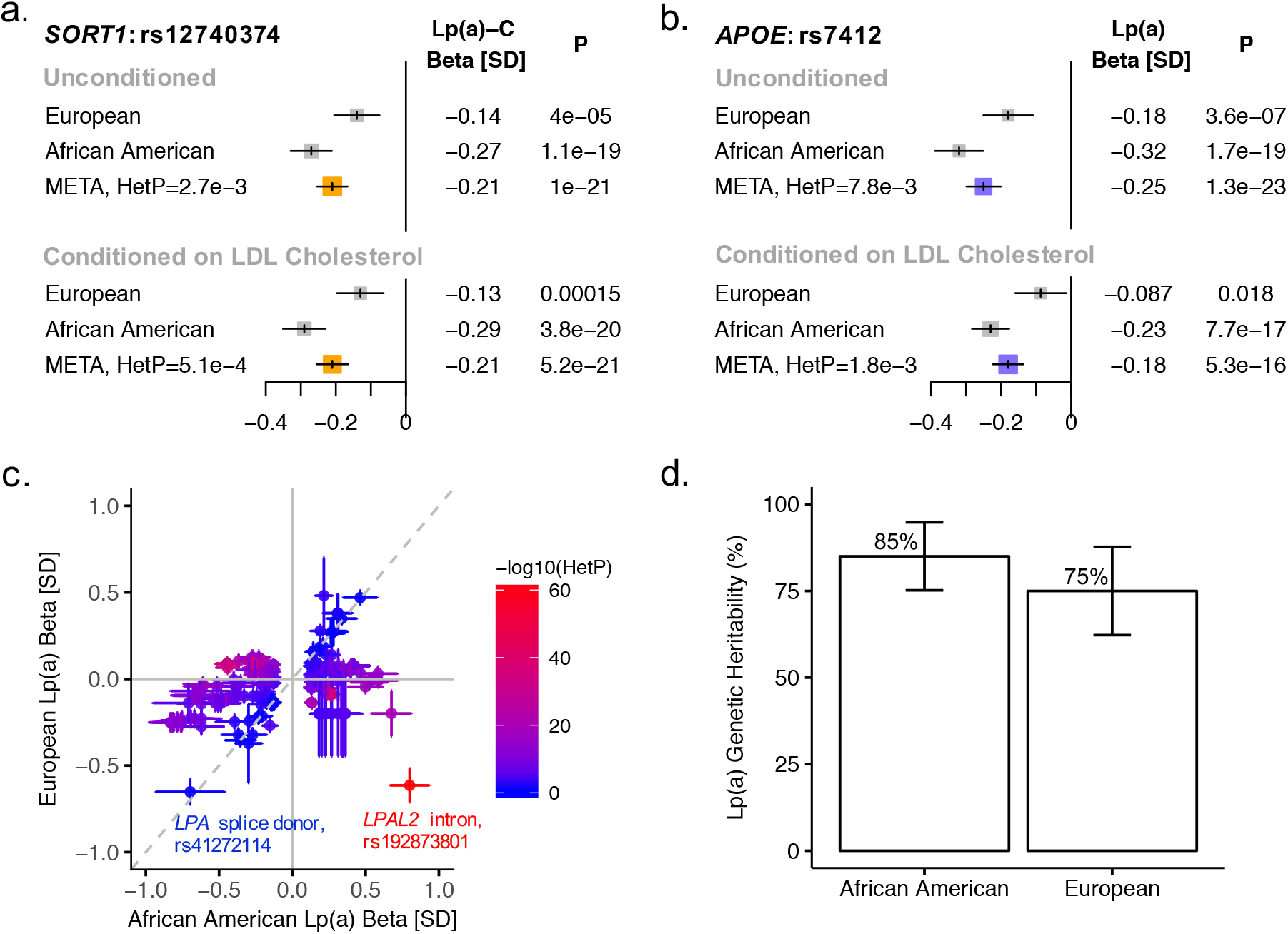
Trans-ethnic *LPA* and non-*LPA* loci associations with lipoprotein(a) phenotypes. In trans-ethnic meta-analysis of single variant results adjusted for KIV2-CN, we observed two associations (*P* < 5×10^−8^) at loci distinct from *LPA* and independent of other conventional lipid measures: *SORT1* for Lp(a)-C and *APOE* for Lp(a). a-b) Associations (Betas in SD and 95% CI) for top variants at the *SORT1* and *APOE* loci are shown by ethnicity. The *SORT1* and *APOE* loci have been previously associated with LDL cholesterol. Thus, associations conditional on LDL cholesterol are also presented. The effect size for *SORT1* is preserved after conditioning on LDL cholesterol while the effect size for *APOE* is slightly reduced but still genome-wide significant. c) Standardized effect estimates for variants at the *LPA* locus *(LPA* TSS +/− 1Mb) attaining *P* < 5x10^−8^ in JHS are shown comparing effects in JHS (African Americans) with FIN (European Americans). Color demonstrates inter-ethnic effect difference as measured by heterogeneity *P*. Similar effects are observed for a known null (splice donor) mutation in *LPA* but strongly diverging effects are observed for a distinct nearby *LPAL2* intronic variant. d) Genetic heritability estimates using variants with MAF > 0.001 for normalized Lp(a) were acquired for African Americans in the whole-genome sequenced JHS cohort and for Europeans in the genotyped and imputed FIN cohort. Here, heritability and 95% CI are shown without adjusting for KIV2-CN. KIV2-CN = kringle IV-2 copy number; HetP = heterogeneity P; Lp(a) = lipoprotein(a); Lp(a)-C = lipoprotein(a) cholesterol; MAF = minor allele frequency; TSS = transcription start site

On average, *LPA* locus genetic variants yielding a 1 SD increase in Lp(a) yield a 0.48 SD increase in Lp(a)-C, similar to the observational correlation between the two phenotypes (**Supplementary Fig. 14**). Iterative conditional analyses at the *LPA* locus showed that, for Lp(a)-C there are 2 (JHS) and 3 (EST) independent genome-wide significant variants, (**Supplementary Table 9a,b**), while for Lp(a) there are 13 (JHS) and 30 (FIN) independent genome-wide significant variants (**Supplementary Table 10a,b**) (**Supplementary Fig. 15a,b**), similar to the number of independent variants from past studies^11,17,30,31^. We replicated Lp(a) associations for two known *LPA* loss-of-function (LOF) alleles^24,25^: splice donor variant rs41272114 (*B* = −0.7 SD, *P* = 8×10^−77^) and splice acceptor variant rs143431368 (*B* = −0.5 SD, *P* = 2×10^−26^), and also discovered a novel LOF variant, a splice acceptor variant in exon 28 only observed African Americans in JHS: rs199583644 (MAF = 0.28%, *B* = −1.5 SD, *P* = 3×10^−13^).

Next, we compared inter-ethnic effects of *LPA* locus variants attaining sub-threshold significance (*P* < 1×10^−4^) in either ethnicity for Lp(a) and Lp(a)-C. Spearman rank correlation of genetic effects between the two ethnicities for Lp(a)-C was 0.38 and for Lp(a) 0.16 (**Supplementary Fig. 16a,b**). Moderately associated (*P* < 1×10^−2^) *LPA* locus variants largely private in African Americans (FIN MAF <0.1%) had larger absolute effects across MAFs compared to such variants observed in both ethnicities (*P* = 3×10^−32^) (**Supplementary Fig. 17a,b**). In comparing betas from genome-wide significant variants in African Americans with betas from the same variants in Europeans (**Fig. 4c**), we found the strongest inter-ethnic heterogeneity *(HetP* = 9.8×10^−64^) at an *LPAL2* intronic variant at the *LPA* locus (rs192873801, MAF 2.8% in JHS and 2.7% in FIN) with strongly divergent effects between the two ethnicities: +0.80 SD in JHS (*P* = 3.8×10^−32^) and −0.61 SD in FIN (*P* = 2.0×10^−35^) **Supplementary Fig. 18**. We noted these variants to be on separate haplotypes for JHS and FIN (**Supplementary Fig. 19**). Notably, the *LPA* loss-of-function variant rs41272114, shows similarly strong effects in both ethnicities (*HetP* > 0.05).

Early family studies in Europeans and Africans have suggested the heritability of Lp(a) to be between 51%−90% ^6–10^. A recent array-based genotyping study in KORA estimated 49%^11^ of variance in Lp(a) from genome-wide heritability analysis of 6,002 Europeans. From WGS, we now estimate genetic heritability in African Americans and Europeans, respectively, to be 85% (SE 5%) and 75% (SE 7%) for Lp(a), and 52% (SE 7%) and 75% (SE 34%) for Lp(a)-C (**Fig. 4d**).

### Common variant association: KIV2-CN modifier analyses

To determine if there are variants that influence the relationship between KIV2-CN and Lp(a)-C or Lp(a) concentrations, we performed variant-by-KIV2-CN interaction analyses at a 4MB window around *LPA*. We identified three independent modifier variants at this locus which influenced the relationship between KIV2-CN and Lp(a)-C (rs13192132, *P* = 1.73×10^−15^, rs1810126, *P* = 6.84×10^−14^, rs1740445, *P* = 6.35×10^−9^) (**Fig. 5**) and were consistent across ethnicities (**Supplementary Table 11, Supplementary Fig. 20a,b**). Sensitivity analyses of interactions was performed to assess for confounding from 1) haplotype effects and 2) single variants tagged through LD^32,33^. All three variants show association with Lp(a)-C individually (*P* < 0.05), but are not correlated with KIV2-CN genotype (Pearson correlation r^2^ < 0.1) (Supplementary Table 12). Furthermore, interaction associations persisted after conditioning on variants independently associated with Lp(a)-C (**Supplementary Table 13**).

**Fig 5.**
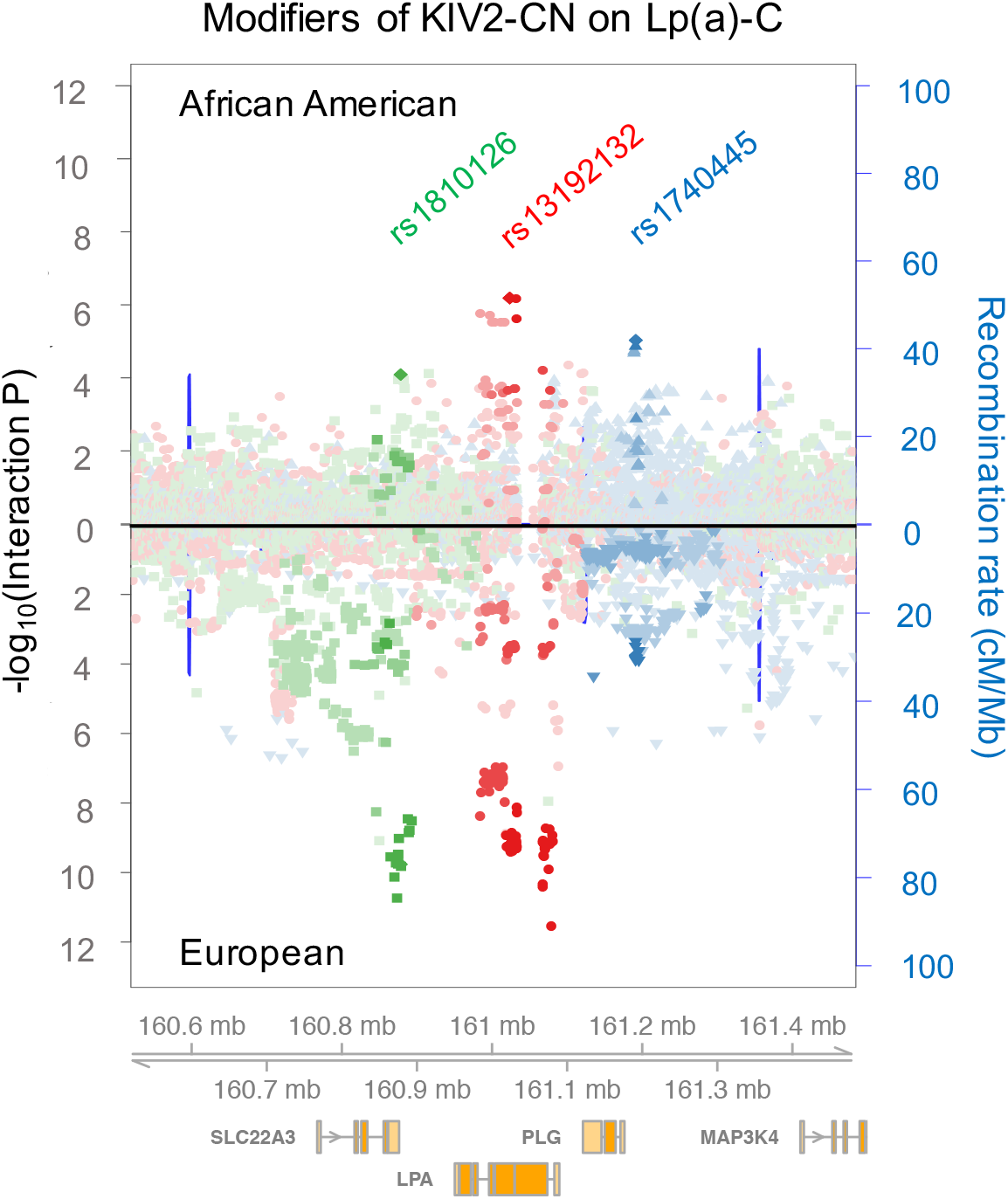
Genetic modifiers of KIV2-CN’s effect on lipoprotein(a) cholesterol. Three independent genetic modifiers of KIV2-CN’s effect on Lp(a)-C were discovered at the *LPA* locus. Regional association plots showing the variant-by-KIV2-CN interaction *P* values of all variants within a 1Mb window of the *LPA* TSS are shown for African Americans (top) and Europeans (bottom), highlighting variants in linkage disequilibrium with rs1810126 (green), rs13192132 (red), and rs1840445 (blue), the top independent genome-wide significant variants (interaction *P* < 5×10^−8^) upon meta-analysis. KIV2-CN = kringle IV-2 copy number;Lp(a)-C = lipoprotein(a) cholesterol; TSS = transcription start site

Genomic context interrogation using adult liver regulatory annotations from the Roadmap Epigenome Project^34^ showed that the top modifier variant in EST, a 3-base deletion, rs4063600 (TAGG>T, *B* = +0.03 SD Lp(a)-C/CN/allele, *P* = 2.96×10^−12^), is in strong LD with rs13192132 (r^2^ = 0.88) and overlies significant H3K4me3 and H3K27ac peaks (*P* < 1×10^−2^) 7,508 bases downstream of the *LPA* transcription start site (TSS) (**Supplementary Fig. 21a**). We additionally performed variant-by-KIV2-CN modifier analyses for Lp(a) using the JHS WGS (**Supplementary Fig. 21b**). Complete lists of cohort-specific, LD-clumped significant variants are provided in **Supplementary Tables 14-17**.

### Rare variant analysis: Coding and non-coding burden tests

Rare and low-frequency disruptive coding variants within *LPA* have been previously associated with Lp(a)^24,25^. Here, we performed two coding rare variant analyses studies (RVAS) aggregating rare (MAF <1%) variants which were 1) LOF or missense deleterious by *in-silico* prediction tools^35^, or 2) non-synonymous, within their respective genes, and performed association with Lp(a)-C, adjusting for KIV2-CN. All analyses were done separately for JHS and EST and meta-analyzed. While no genes reached significance in either analysis after accounting for multiple-hypothesis testing, we observed suggestive evidence for *LPA* in both coding RVAS tests (*P* = 7×10^−4^ for LOF and missense deleterious mutations, 1×10^−4^ for non-synonymous mutations) (**Supplementary Tables 18-19, Supplementary Fig. 22a,b**).

We also interrogated whether there was evidence of rare, non-coding variants aggregated within regulatory sequences uniquely detected by WGS that influence Lp(a)-C. We performed three non-coding RVAS using the variant groupings described in the Methods along with Roadmap epigenome data^34^ from adult liver, the main tissue where *LPA* is expressed (**Supplementary Fig. 23, Supplementary Fig. 24**). The only genome-wide significant association was for an intron of *SLC22A3* at 6:160851000-160854000 with Lp(a)-C (*P* = 4.5×10^−8^) (**Supplementary Tables 20-25**). Similarly, rare variants in a putative regulatory domain of *SLC22A3* were recently shown to be associated with Lp(a) in a sliding window analysis using low-coverage whole genomes^36^. However, we found that conditioning on *LPA’s* KIV2-CN, 128 kb away, mitigated the observed association (*P* = 4.3×10^−3^, **Supplementary Tables 20-21**). Upon conditioning on KIV2-CN, while no sliding windows reached statistical significance, the top window was 6:160,939,500-160,942,500 (*P* = 1.6×10^−4^), 13kb downstream of the *LPA* transcription end site and overlapping three annotated ORegAnno^37^ CTCF binding sites (**Fig. 6**).

**Fig 6.**
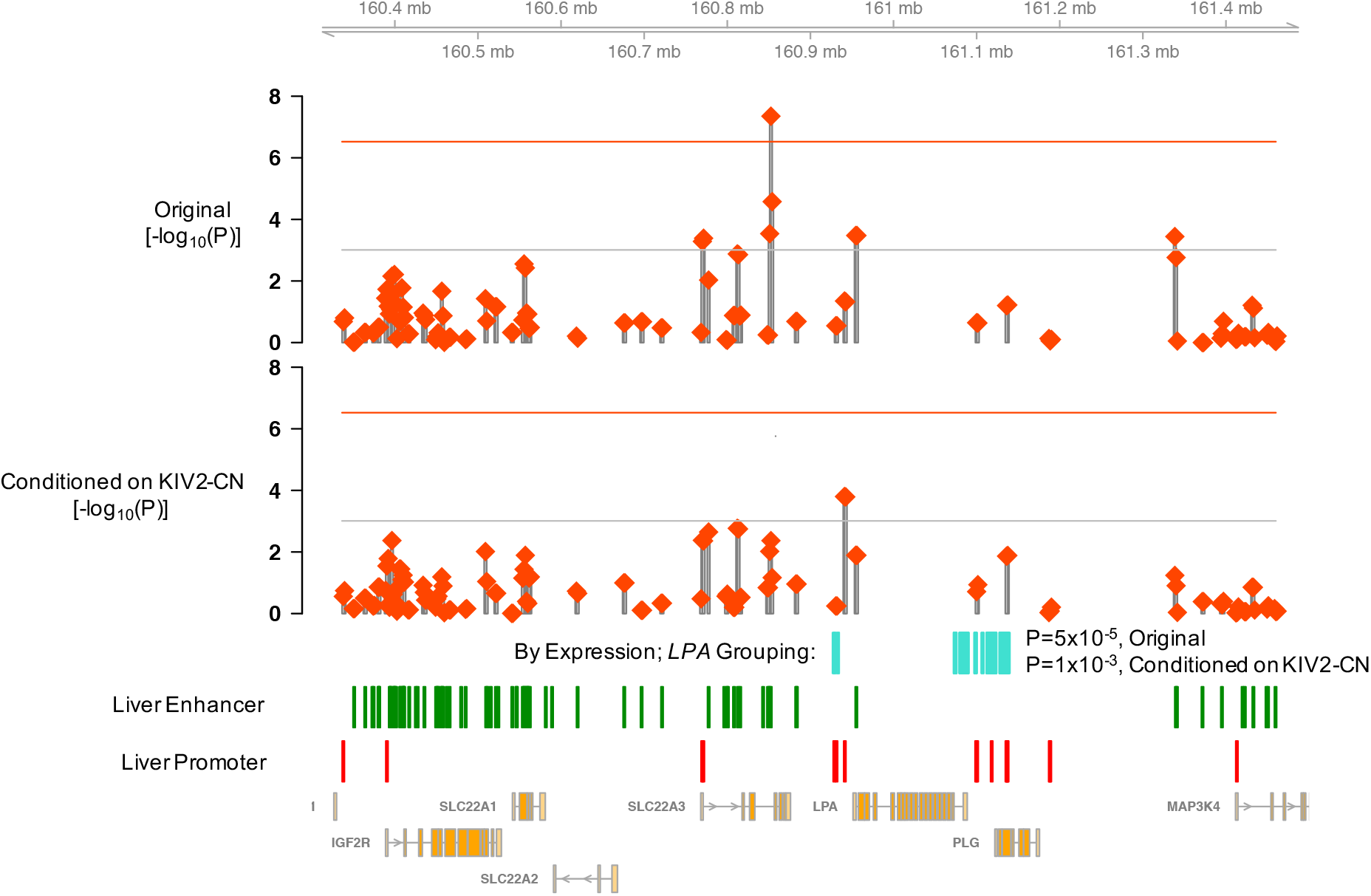
Rare variant non-coding burden analyses. A schematic of rare variant association results from (1) aggregating rare variants in adult liver enhancers or promoters and strong DHS (*P* < 10^−10^) within 3kb sliding windows, and (2) aggregating rare variants in liver enhancers grouped to *LPA* via the “By expression” in-silico prediction method. In the top two panels, each red diamond represents the meta-analyzed mixed-model SKAT P-value with Lp(a)-C of rare (MAF < 1%), non-coding variants overlapping liver enhancer or promoter annotations in strong DHS (P(DHS) < 1e-10) grouped in a 3kb window, before adjusting for KIV2-CN (top, “Original”) and after adjusting for KIV2-CN (bottom, “Conditioned on KIV2-CN”). The horizontal red lines denote the genome-wide Bonferroni significance threshold given the number of unique windows analyzed. The horizontal gray lines denote the Bonferroni significance threshold within this 1MB region around *LPA*. The regions incorporated into the “By Expression” grouping to *LPA* are shown in aqua, along with the respective associations of rare non-coding variants in these regions before and after conditioning on KIV2-CN. Annotated adult liver enhancers (green bars) and promoters (red bars) overlapping strong DHS are included above protein-coding genes from ENSEMBLE. DHS = DNAse hypersensitivity sites; Lp(a)-C = lipoprotein(a) cholesterol; MAF = minor allele frequency; SKAT = Sequence Kernal Association Test

Interrogation of rare enhancer variants predicted to influence *LPA* expression in liver^38^ showed nominal evidence of association with Lp(a)-C before (*P* = 5×10^−5^) and after (*P* = 1×10^−3^) conditioning on KIV2-CN (**Fig. 6, Supplementary Fig. 25**). However, other putative gene-linked rare enhancer variants at the *LPA* locus, including the aforementioned *SLC22A3* (**Supplementary Fig. 26**), also demonstrate nominal associations, highlighting current challenges in both mapping associated regulatory elements to causal genes through *in silico* approaches and discerning the relative impacts of potentially pleiotropic regulatory elements.

### Mendelian randomization

Genetic variation at the *LPA* locus is an optimal instrument for Mendelian randomization (MR) as it strongly and specifically influences circulating Lp(a) levels. Past studies have performed Lp(a) MR across clinical and metabolic traits using genetic risk scores comprised of between 1-18 variants^14,39,40^. Here, we performed MR using three different genetic instruments per cohort to distinguish variant classes influencing Lp(a) phenotypes: 1) an expanded genetic risk score, “GRS,” comprised of the sum of the KIV2-CN-adjusted variant effects from LD-pruned variants in a ~4MB window around *LPA* with sub-threshold significance (*P* < 1×10^−4^); 2) a “KIV2-CN” score using the directly genotyped or imputed KIV2-CN; and 3) a combined “GRS+KIV2-CN” score combining scores from (1) and (2). Each genetic instrument was normalized such that 1 unit increase in the score was equal to 1SD increase in Lp(a) (or Lp(a)-C). In African Americans, 235 variants were used towards the Lp(a) GRS and 39 towards the Lp(a)-C GRS (**Supplementary Table 26**). In Europeans, 399 variants were used towards the Lp(a) GRS and 49 towards the Lp(a)-C GRS (**Supplementary Table 27-28**). The GRS+KIV2-CN score explains 45-49% of Lp(a) variance and 20% of Lp(a)-C variance (**Supplementary Fig 27, Supplementary Table 29**).

Association of GRS+KIV2-CN with 10 incident clinical events from the FIN imputation dataset (N=27,344) (**Fig. 7a, Supplementary Table 30**) demonstrated anticipated associations for incident cardiovascular diseases (HR 1.18/Lp(a) SD, *P* = 1×10^−5^), comprising incident myocardial infarction (HR 1.23/Lp(a) SD, *P* = 8×10^−4^), coronary heart disease (CHD) (HR 1.25/Lp(a) SD, *P* = 7×10^−7^), and stroke (HR 1.27/Lp(a) SD, *P* = 1×10^−3^). For given effect on Lp(a), the GRS had a larger effect on incident cardiovascular risk (HR 1.30/Lp(a) SD, *P* = 6×10^−8^) than KIV2-CN (HR 1.03/Lp(a) SD, *P* = 0.17). While the KIV2-CN score alone was not as strongly associated with cardiovascular outcomes (*P* > 0.05), its estimated effect with incident MI (HR = 1.16) was similar to recent estimations in a MI case-control analysis^14^. Thus, power for MR using the KIV2-CN instrument may be hindered due to a limited number of incident MI cases and modest effect conferred by KIV2-CN. These results suggest that knowledge of *LPA* variant class genotypes may provide additional information on cardiovascular risk beyond circulating Lp(a) levels.

**Fig 7.**
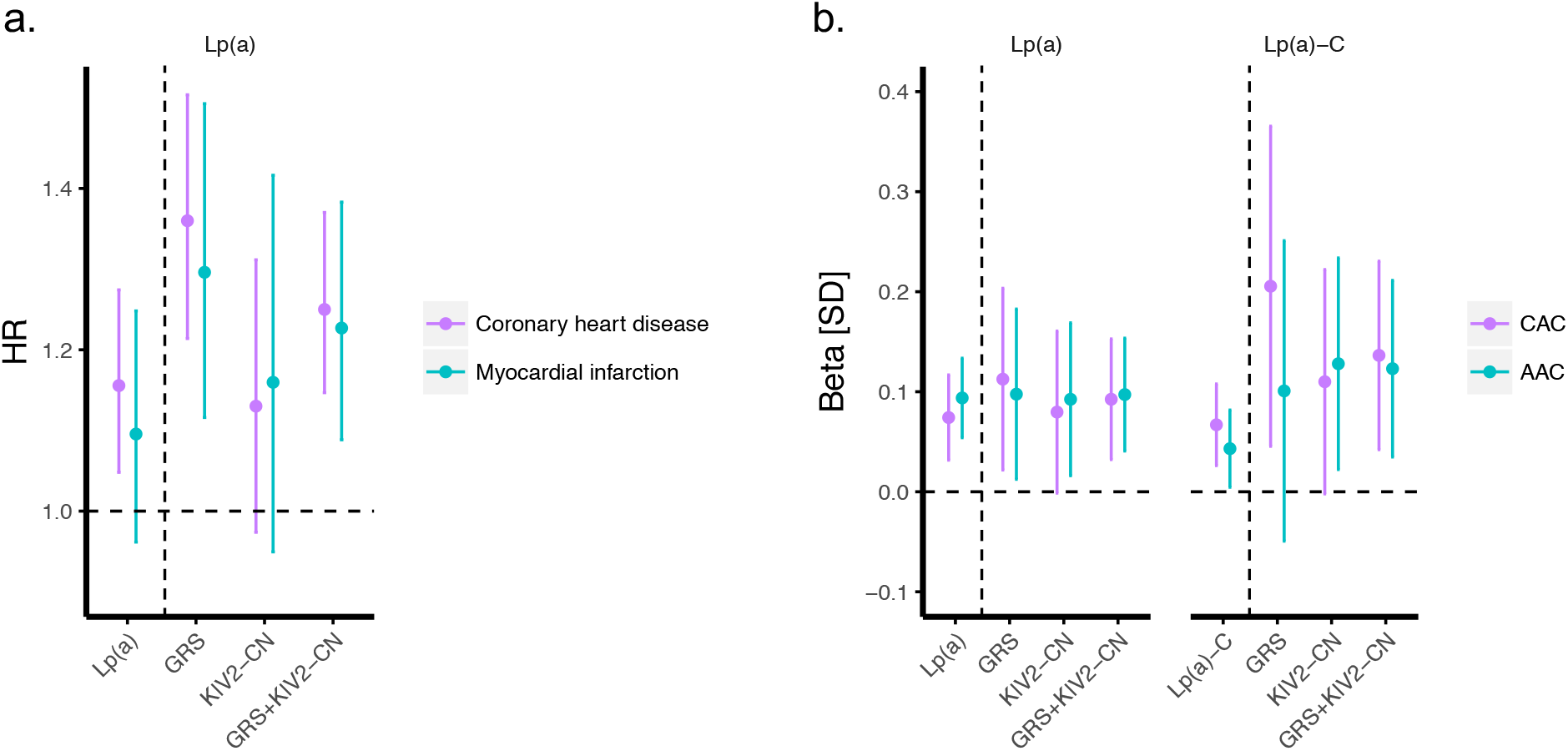
Association of *LPA* variant classes with incident clinical events and subclinical measures. Mendelian randomization was performed using three genetic instruments: a weighted genetic risk score using variants conditioned on KIV2-CN at a 4Mb window around *LPA* (GRS), a KIV2-CN score, and a combined GRS+KIV2-CN score, and compared to the observational effects. The genetic instruments were all normalized such that 1 unit increase in the score is equal to 1SD increase in Lp(a) or Lp(a)-C. a) Associations (HR and 95% CI) of incident coronary heart disease (1,056 cases; 21,207 controls) and myocardial infarction (580 cases; 21,377 controls) with the Lp(a) measurement and with genetic instruments among the genotyped and imputed FIN individuals (exact values in **Supplementary Table 30**). b) Associations (Beta and 95% CI) of Lp(a) and Lp(a)-C measurements and respective genetic instruments with standardized markers of subclinical atherosclerosis (CAC and AAC) among 1,701 whole-genome sequenced JHS participants (exact values in **Supplementary Table 31**). These data indicate that 1) a comprehensive Lp(a) genetic instrument (GRS+KIV2-CN) provides improved risk assessment compared to the Lp(a) phenotype, and 2) further stratifying this comprehensive instrument into separate Lp(a) variant classes provides additional risk stratification in that genomic sequence variants independent of KIV2-CN (i.e.: GRS) have a stronger influence on clinical atherosclerosis compared to KIV2-CN. AAC = Abdominal aortic calcium; CAC = coronary artery calcium; CI = confidence interval; FIN = FINRISK; GRS = genetic risk score; HR = hazard ratio; JHS = Jackson Heart Study; KIV2-CN = kringle IV-2 copy number; Lp(a) = lipoprotein(a); Lp(a)-C = lipoprotein(a) cholesterol

To determine whether *LPA* genomic variants influence the accumulation of subclinical cardiovascular atherosclerosis, we associated both the Lp(a) and Lp(a)-C genetic instruments with computed tomography-derived measures of atherosclerosis in the coronary arteries (CAC) and abdominal aorta (AAC) in 1,701 African Americans from JHS without prevalent clinical cardiovascular disease (**Supplementary Table 31, Fig. 7b**). Here, the comprehensive (GRS+KIV2-CN) genetic instruments for both Lp(a) and Lp(a)-C demonstrated association with subclinical atherosclerosis with similar standardized effects for both CAC and AAC: Lp(a) (CAC: *B* = 0.097, *P* = 7.6×10^−4^; AAC: *B* = 0.092, *P* = 2.7×10^−3^), and Lp(a)-C (CAC: *B* = 0.14, *P* = 4.6×10^−3^; AAC: *B* = 0.12, *P* = 6.4×10^−3^). Notably, this is the first known demonstration of *LPA* genomic variants affecting atherosclerotic risk in African Americans. A prior study of African Americans from the Dallas Heart Study found no association between sub-clinical measures of atherosclerosis, such as coronary calcium, and Lp(a) phenotype^41^. Compared to prior work, our power is optimized with larger sample size and genetic instrument for causal inference.

## DISCUSSION

We characterized the genetic architecture of Lp(a) and Lp(a)-C using deep coverage WGS in 8,392 Europeans and African Americans across allele frequencies and classes. While we observe that Lp(a) is highly heritable in Europeans and African Americans, distinct and common genetic determinants influence concentrations. Using a comprehensive genetic instrument that separately imputes apo(a) isoform, we show that knowledge of *LPA* genotypes can better inform incident cardiovascular disease risk prediction than just knowledge of Lp(a) biomarker level.

These observations permit several conclusions. First, through whole-genome sequencing and imputation, we observe substantial genetic heritability of Lp(a) −85% (SE 5%) in African Americans and 75% (SE 6%) in Europeans. We leverage this observation to systematically dissect the heritable components of Lp(a) across two ethnicities. Through single variant analysis, we find a novel locus for Lp(a)-C, *SORT1*, whereby the top variant (rs12740374) reduces plasma Lp(a)-C concentrations in both ethnicities and is independent of LDL cholesterol levels, thereby providing evidence for the sortilin receptor as a novel component in Lp(a)-C metabolism. Through genetic modifier analysis, we find evidence of three loci which affect the relationship between KIV2-CN and Lp(a)-C similarly across both ethnicities. We replicate evidence supporting rare coding variation at *LPA* influencing Lp(a); however, observed associations of aggregates of rare non-coding variation appeared to be largely explained by LPA structural variation, namely KIV2-CN.

Second, we observed high heritability in diverse ethnicities despite notable inter-ethnic differences in circulating biomarker concentrations. Upon finding that similar Lp(a) effect sizes are conferred per KIV2 copy in African Americans and Europeans, we delved further into KIV2-independent effects conferred by variants at the *LPA* locus. Among distinct sequence variation, we notably observed an *LPAL2* intronic variant with significant yet opposing effects in each ethnicity, likely indicating influences from haplotype structure or gene-environment interactions. Altogether, *LPA* locus variants largely private to African Americans (FIN MAF < 0.1%) confer significantly greater absolute effect on standardized Lp(a) levels than variants observed in both ethnicities.

Third, WGS enables the detection of relevant genomic variants for Lp(a) which cannot be detected via WES or genotyping arrays. Furthermore, knowledge of such variants, given differential effects on circulating Lp(a) and differential effects on incident cardiovascular events, provides additional information regarding cardiovascular disease risk beyond circulating Lp(a).

It should be noted that several limitations to this work exist. First, we estimate total KIV2-CN, but individuals may have different KIV2-CN alleles on each chromosome^42^. Our CNV analysis of next-generation sequencing data relies on aggregate depth of coverage for genotyping, precluding our ability to determine allelic KIV2-CN. However, despite this, sensitivity analyses suggest that the sum of KIV2-CN alleles may similarly associate with Lp(a) across varied KIV2-CN allele combinations. Additionally, the strongest SNP in our KIV2-CN imputation model is rs10455872, whose association with KIV2-CN has been well-described previously^17^, and our KIV2-CN estimate is robustly associated with Lp(a) phenotypes as expected. Second, we only assess one non-European cohort; however, it has been observed that there are distinct Lp(a) distributions in other ethnicities which may uncover additional loci and sources of genetic heterogeneity. Third, while *in silico* prediction tools for non-coding regions identify putative regulatory sequence, they are limited in their ability to 1) determine disruptive mutations, and 2) link regulatory regions to genes.

In summary, we characterize the shared and unique genetic determinants of Lp(a) using whole genome sequences in African Americans and Europeans. Additional knowledge of the complement of these determinants better informs cardiovascular disease risk prediction than biomarker alone.

Uncategorized References

## ACKNOWLEDGEMENTS

Whole genome sequencing (WGS) for the Trans-Omics in Precision Medicine (TOPMed) program was supported by the National Heart, Lung and Blood Institute (NHLBI). WGS for “NHLBI TOPMed: The Jackson Heart Study” (phs000964.v1.p1) was performed at the University of Washington Northwest Genomics Center (HHSN268201100037C). Centralized read mapping and genotype calling, along with variant quality metrics and filtering were provided by the TOPMed Informatics Research Center (3R01HL-117626-02S1). The Jackson Heart Study (JHS) is supported and conducted in collaboration with Jackson State University (HHSN268201300049C and HHSN268201300050C), Tougaloo College (HHSN268201300048C), and the University of Mississippi Medical Center (HHSN268201300046C and HHSN268201300047C) contracts from the National Heart, Lung, and Blood Institute (NHLBI) and the National Institute for Minority Health and Health Disparities (NIMHD). Phenotype harmonization, data management, sample-identity QC, and general study coordination, were provided by the TOPMed Data Coordinating Center (3R01HL-120393-02S1). We gratefully acknowledge the studies and participants who provided biological samples and data for TOPMed, Estonia, and FINRISK cohorts. The authors also wish to thank the staffs and participants of the JHS, Estonia, and FINRISK cohorts. S.K. is supported by an Ofer and Shelly Nemirovsky Research Scholar Award from Massachusetts General Hospital, RO1 HL127564 from the National Heart, Lung, and Blood Institute, and UM HG008895 from the National Human Genome Research Institute. T.E. and A.M. are funded by Estonian Research Council Grant IUT20-60 and PUT1660, EU H2020 grant 692145, and European Union through the European Regional Development Fund (Project No. 2014-2020.4.01.15-0012) GENTRANSMED. J.G.W. is supported by U54GM115428 from the National Institute of General Medical Sciences. I.S. is supported by the Academy of Finland (298149). V.S is supported by the Finnish Foundation for Cardiovascular Research. S.M.Z is supported by the Paul & Daisy Soros Fellowship. The views expressed in this manuscript are those of the authors and do not necessarily represent the views of the National Heart, Lung, and Blood Institute; the National Institutes of Health; or the U.S. Department of Health and Human Services.

## AUTHOR CONTRIBUTIONS

Concept and design: S.M.Z., P.N., A.G., I.S., S.K., Analysis: S.M.Z., S.R., R.E.H., Interpretation of data: S.M.Z., S.R., R.E.H., P.N., S.K., S.R., R.E.H., M.A., M.J.D., I.S., A.G., Drafting of the manuscript: S.M.Z., P.N., A.G., S.K., Revision of the manuscript: S.M.Z., P.N., A.G., S.K., I.S., R.E.H., M.A., V.S., J.G.W., S.R., Hail software support: J.B., T.P., C.S., Other technical support: J.Ernst, J.Engreitz, Administrative or material support: A.C., A.M., V.S., M.K., M.J.D, J.G.W., B.M.N., S.M., I.S., T.E., S.R., S.K., P.N..

## METHODS

### Study participants

Please refer to **Supplementary Text** for study participant details.

### Whole genome sequencing and variant calling

Sequencing was performed at one of two sequencing centers, with all members within a cohort sequenced at the same center. The JHS WGS individuals were sequenced at University of Washington Northwest Genomics Center (Seattle, WA) as part of the as a part of the Phase 1 NIH/NHLBI Trans-Omics for Precision Medicine (TOPMed) program. The Finnish and Estonian WGS individuals were sequenced at the Broad Institute of Harvard and MIT (Cambridge, MA). Target coverage was >30X for JHS (mean attained 37.1), >20X for EST (mean attained 30.4), and >20X for FIN (mean attained 29.8).

TOPMED phase 1 BAM files were harmonized by the TOPMed Informatics Research Center (Center for Statistical Genetics, University of Michigan, Hyun Min Kang, Tom Blackwell and Goncalo Abecasis). In brief, sequence data were received from each sequencing center in the form of bam files mapped to the 1000 Genomes hs37d5 build 37 decoy reference sequence. Processing was coordinated and managed by the ‘GotCloud’ processing pipeline^43^. Samples with DNA contamination > 3% (estimated using verifyBamId software^44^) and <95% of the genome covered at least 10x were filtered out. The JHS WGS used for analysis are from the “freeze 3a” genotype callsets of the variant calling pipeline performed using the software tools in the following repository: https://github.com/statgen/topmed_freeze3_calling, with variant detection performed by vt discover2 software tool^45^.

WGS for FINRISK and the Estonian Biobank were performed using the Illumina HiSeqX platform at the Broad Institute of Harvard and MIT (Cambridge, MA). Libraries were normalized to 1.7nM, constructed, and sequenced on the Illumina HiSeqX with the use of 151-bp paired-end reads for WGS and output was processed by Picard to generate aligned BAM files (to hg19) ^46,47^. Variants were discovered using the Geome Analysis Tookit (GATK) v3 HaplotypeCaller according to Best Practices^48^. Finland and Estonia WGS samples were jointly called.

### Whole genome sequence quality control

#### Sample quality control

The following three approaches were used by the TOPMed Genetic Analysis Center to identify and resolve sample identity issues in JHS: (1) concordance between annotated sex and biological sex inferred from the WGS data, (2) concordance between prior SNP array genotypes and WGS-derived genotypes, and (3) comparisons of observed and expected relatedness from pedigrees.

Additional measures for quality control of JHS, Finland, and Estonia were performed using the Hail software package (https://github.com/hail-is/hail)^49^. Samples were filtered by contamination (>3.0% for JHS, >5.0% for Finland and Estonia), chimeras >5%, GC dropout >4, raw coverage (<30X for JHS, <19X for Finland and Estonia), and indeterminant genotypic sex or genotypic/phenotypic sex mismatch (**Supplementary Table 1**).

#### Genotype and Variant quality control

The variant filtering in JHS was performed by (1) first calculating Mendelian consistency scores using known familial relatedness and duplicates, and (2) training SVM classifier between the known variant sites (positive labels) and the Mendelian inconsistent variants (negative labels). Two additional hard filters were applied: (1) Excess heterozygosity filter (EXHET), if the Hardy-Weinberg disequilbrium p-value was less than 1×10^−6^ in the direction of excess heterozygosity; (2) Mendelian discordance filter (DISC), with 3 or more Mendelian inconsistencies or duplicate discordances observed from the samples. Genotypes with a depth < 10 were excluded, prior to filtering variants with > 5% missingness.

Variants for Finland and Estonia were initially filtered by GATK Variant Quality Score Recalibration. Additionally, genotypes with GQ<20, DP<10 or >200, and poor allele balance (homozygous with <0.90 supportive reads or heterozygous with <0.20 supportive reads) were removed. Variants within low complexity regions were removed across all samples.^50^ Variants with > 20% missing calls, quality by depth <2 (SNPs) or <3 (indels), InbreedingCoeff <−0.3, and pHWE <1×10^−9^ were filtered out.

### Finnish imputation and quality control

The imputation of the FINRISK samples^51^ was done utilizing population specific reference panel of 2,690 high-coverage whole-genome and 5,093 high-coverage whole-exome sequences with IMPUTE2^52^ that allows the usage of two panels at the same time. Before phasing and imputation, the data was QCed using following criteria: exclude samples with obscure sex, missingness (>5%), excess heterozygosity (+-4sd), non-European ancestry and SNPs with low call-rate (>2% missing), low HWE P-value (<1e-6), minor allele count (MAC) < 3 (in case Zcalled^53^) or MAC < 10 (if only called using Illumina GenCall). The haplotypic phase was determined using SHAPEIT2.0^54^ prior to imputation. The FINRISK samples have been genotyped using multiple different genotyping chips, for which the QC, phasing and imputation was done in multiple chip-wise batches.

### Lipid phenotypes

#### Lp(a) and Lp(a)-C

Serum Lp(a)-C was measured in both EST and JHS via density gradient ultracentrifugation (Vertical Auto Profile [VAP], Atherotech).

Lp(a) was measured in JHS using a Diasorin nephelometric assay on a Roche Cobas FARA analyzer (Roche Diagnostics Corporation, Indianapolis, IN, USA), which measures Lp(a) mass by immunoprecipitin analysis using the SPQTM Antibody Reagent System of DiaSorin (DiaSorin Inc., Stillwater, MN 55082-0285). Turbidity produced by the antigen-antibody complexes was measured using the Roche Modular P Chemistry Analyzer. In FIN, Lp(a) was measured from serum stored at −70°C using a commercially available latex immunoassay on an Architect c8000 system (Quantia Lp(a), Abbott Diagnostics).

Lp(a)-C and Lp(a) were inverse-rank normalized separately by cohort for analysis.

#### Conventional lipids

Conventional lipoprotein cholesterols (HDL, LDL, TG, Total Cholesterol) and proteins (ApoB, ApoAI) were measured in EST and JHS by the VAP assay (where LDL refers to directly measured LDL, and not calculated). In FIN, these lipoproteins were measured via NMR as described in the Mendelian randomization methods below. In FIN, LDL cholesterol was either calculated by the Friedwald equation when triglycerides were <400 mg/dl or directly measured. Given the average effect of statins, when statins were present, total cholesterol was adjusted by dividing by 0.8 and LDL cholesterol by dividing by 0.7, as previously done^55^. All lipids were inverse-rank normalized separately by cohort in analysis.

### KIV2-CN estimation from WGS data

Genome STRiP^21^ version 2.00.1710 was used to estimate KIV2-CN in the *LPA* gene.

Specifically, we ran Genome STRiP read-depth genotyping on the hg19 interval 6:161032614-161067851 using the following custom settings to capture an aggregate read depth signal over every base position: -P depth.minimumMappingQuality:0, without specifying any of the usual genome masks.

After genotyping, we estimated the number of KIV2 protein domains from the raw copy number estimate by dividing the VCF genotype field CNF by the info field GSM1 and then estimating the KIV2 copy number by

> KIV2-CN = (CNF / GSM1) * 6.354 – 0.708

where 6.354 is derived from the number of full copies of the repeating unit represented on the hg19 reference genome and -0.708 is to adjust to the KIV2 units as visualized in **Supplementary Fig. 6A**, removing the outermost flanking exons that are part of the KIV1 and KIV3 (which are picked up in Genome STRiP due to their homology with the exons within the KIV2 domain).

### Evaluation of KIV2-CN precision

To evaluate the precision of our measurements of KIV2 copy number, we utilized 123 pairs of siblings from JHS that were confidently IBD2 (identical by descent on both haplotypes) at the *LPA* locus. To identify these sibling pairs, we interrogated the hg19 interval 6:160,450,001-161,590,000 (0.5 Mb upstream and downstream of the *LPA* gene) and computed the concordance of SNP genotypes in this interval between all sequenced sibling pairs. We classified all sibling pairs with less than 1% genotype discordance as confidently IBD2 at the *LPA* locus and compared IBD2 sibling KIV2-CNs.

### KIV2-CN Imputation

We split the FIN WGS into one training dataset comprised of two thirds of the samples (1477 samples) and one validation dataset (738 samples), and used the least absolute shrinkage and selection operator (LASSO), a machine-learning regression analysis method, using variants in a 4MB window around *LPA* imputed with high-quality (imputation quality > 0.8) and MAF > 0.001 in the FIN dataset. After applying 10-fold cross validation to find the optimal lambda (degree of shrinkage), the LASSO model selected 61 variants which minimized the mean squared error (**Supplemental Fig. 6A**). These 61 variants were also used in a random forest model, reiterating the exponential decay of mean-squared error as the number of variants in the model reaches 61 (**Supplemental Fig. 6B**), and finding the relative importance of each variant in the model.

### Principle component analysis (PCA)

To visualize PCs across all 3 cohorts against each other, a panel of approximately 16,000 ancestry informative markers^56^ (AIMs) identified across six continental populations^57^ was chosen to derive principal components (PCs) of ancestry for all samples that passed quality control. Principal component analysis was performed using EIGENSTRAT, using suggested quality control criteria^58^ (**Supplementary Fig. 3**). Separately, within-cohort PCA was performed for use as covariates in analysis.

### Variant annotation

Variants were annotated with Hail^49^ using annotations from Ensembl’s Variant Effect Predictor (VEP), ascribing the most severe, canonical consequence and gene to each variant^59^. For noncoding regions in adult liver cells (E066), we used the Reg2Map HoneyBadger2-intersect^34^ at strong (*P* < 1×10^−10^) DNase I hypersensitive regions (https://personal.broadinstitute.org/meuleman/reg2map/HoneyBadger2-intersectrelease/). Variants overlapping putative enhancers and promoters from the 25-state chromatin model^34^ at this link were annotated and used in the single variant results annotations (**Supplementary Tables 7-8**) as well as grouping rare variants in the “sliding window” and “by distance” non-coding rare variant studies.

### Single variant association

Single variant analysis for EST and JHS WGS was performed using Hail’s linear mixed model regression^49^ for associating each variant site with inverse normal transformed Lp(a) and Lp(a)-C within each cohort. All analyses were adjusted for KIV2-CN, age, sex, and an empirically derived kinship matrix to account for both familial and more distant relatedness^60^. To create the kinship matrix, regions of high-complexity known to have high LD were removed (as in the EPACTS make-kin --remove-complex flag); these regions included: 5:44000000-52000000, 6:24000000-36000000, 8:8000000-12000000, 11:42000000-58000000, and 17:40000000-43000000. Ten-fold random down-sampling of variants was performed to further reduce variant counts for fast processing-time.

For the FIN imputation dataset, single variant analysis was performed using SNPTEST (v2.5.2), using KIV2-CN, age, sex, fasting > 10hr, and adding PC1-10 as covariates to account for population structure due to absence of kinship matrix.

To ensure robust results, we only performed single variant analysis for variants with a MAF > 0.001 within either cohort. Summary statistics for JHS and FIN for Lp(a) and JHS and EST for Lp(a)-C, for the corresponding inverse-rank normalized phenotypes, were meta-analyzed across cohorts using METAL^61^, while also calculating heterogeneity statistics. Statistical significance alpha of 5×10^−8^ was used for these analyses.

Additionally, for the *LPA* locus, iterative conditional association analysis was performed by cohort. Iterative conditioning was performed until *P* > 5×10^−8^ was attained.

### Heritability analyses

Heritability analyses in EST WGS (for Lp(a)-C) and JHS WGS (for both Lp(a) and Lp(a)-C) were performed using Hail’s linear mixed model regression heritability estimate^49^, described here https://hail.is/hail/hail.VariantDataset.html?highlight=lmm#hail.VariantDataset.lmmreg. Several filters were applied before variants were used in the kinship matrix. First, genome-wide variants underwent two-fold LD pruning as previously described via BOLT-REML^62^, using variants with MAF > 0.001 and missingness < 1% with maximum LD r^2^ = 0.9 (PLINK^63^ commands used: -- maf 0.001 --geno 0.01 --indep-pairwise 50 5 0.9). Regions of high-complexity were removed as previously described for single variant analysis. Ten-fold random down-sampling of variants was performed to further reduce variant counts for feasible analysis processing-time. For the heritability estimates provided, 6,370,696 variants were used towards the kinship matrix in EST Lp(a)-C analysis, 1,897,407 variants in JHS Lp(a)-C analysis, and 1,894,291 variants in the JHS Lp(a) analysis. Baseline covariates used in the model, performed separately by cohort, included age, sex, fasting > 10h, and for EST, sequencing batch. A separate heritability estimate was also derived additionally conditioning on KIV2-CN.

For the FIN imputation dataset, variants were similarly limited, filtering to variants with MAF > 0.001, imputation quality > 0.8, and applying two-fold LD-pruning and removal of complex regions as described above (though the ten-fold down-sampling was not applied to keep the variant count on the same order of magnitude as in the WGS heritability analyses). A total of 3,088,864 variants were used towards heritability analysis, which was performed using BOLT-REML. Covariates used in the analysis included age, sex, fasting > 10hr, and PC1-10. A separate heritability estimate was also derived additionally conditioning on KIV2-CN. For Lp(a), heritability analysis additionally conditioning on both KIV2-CN and the KIV2-CN-independent GRS using in Mendelian randomization was performed. BOLT-REML was also applied towards the Lp(a) heritability analysis in JHS, arriving at the same heritability estimates as Hail (data not shown).

### KIV2-CN modifier analysis

Variant-by-KIV2-CN interaction analysis in the WGS was performed at a ~4MB window (6:158532140-162664257) around *LPA* to identify variants which modify the relationship between directly genotyped KIV2-CN and Lp(a)-C (for EST and JHS) and Lp(a) (for JHS only). Variants with minor allele count > 20 (by cohort) were included in analyses. The following interaction model was performed: Lp(a)-C ~ KIV2-CN + Variant + KIV2-CN x Variant + covariates Where the interaction effect and p-value corresponds to the term: “KIV2-CN x Variant”. Cohort-specific analyses were performed and for Lp(a)-C, EST and JHS interaction results were meta-analyzed using METAL^61^. Using the full interaction results, three top modifier variants were identified (rs13192132, rs1810126, and rs1740445) that were genome-wide significant upon meta-analysis (*P* < 5×10^−8^), in linkage equilibrium (r^2^ < 0.1) across both ethnic backgrounds, and had replicating interaction effect directions in both ethnicities. To determine the cohort-specific Bonferroni significance threshold, LD clumping was performed on the full interaction results separately by cohort using the following PLINK^63^ flags: --clump-kb 500 --clump-p1 1 --clump-p2 1 --clump-r2 0.25. In JHS, 1373 LD-pruned variants were identified, leading to a significance threshold of *P* = 3.64×10^−5^. In EST, 566 LD-pruned variants were identified, leading to a significance threshold of *P* = 8.83×10^−5^. Clumped variants with interaction p-values surpassing the Bonferroni threshold are provided by cohort and phenotype in **Supplementary Table 14**. For each of these variants, the statistics for the KIV2-CN term, the Variant term, and the KIV2-CN x Variant term from the interaction model, and the variant associations with KIV2-CN and Lp(a)-C (or Lp(a)) are provided in **Supplementary Tables 15–17** (where each Supplementary Table reflects a different cohort and phenotype combination). Overlap with methylation and acetylation marks was visualized using data from Roadmap for E066 adult liver cells at http://egg2.wustl.edu/roadmap/data/byFileType/alignments/consolidated/. Liver ATAC-seq data was downloaded from the ENCODE data portal (accession ENCFF893CSN). FASTQ files were adapter-trimmed and aligned to hg19 with bowtie2, and duplicates reads and reads with MAPQ < 30 were removed.

Previous publications of variant-by-variant interactions have recommended performing sensitivity analyses to ensure significant interactions identified are not 1) due to the variants being in LD on the same haplotype and 2) mitigated by a separate third variant which explains the entire association^32,33^. In particular, the most recent study by Fish et al. ^28^ recommended that variant-by-variant interactions be performed using un-correlated variants (LD r^2^ < 0.6). Thus, we checked the correlation of each of the three top identified variants with KIV2-CN by cohort (**Supplementary Table 12**), finding that these variants are indeed not correlated with KIV2-CN (Pearson correlation r^2^ < 0.1). Furthermore, variants not associated (*P* > 0.05) with the phenotype are suggested to be removed, under the hypothesis that they may represent weak marginal effects from a true underlying interaction. Indeed, our three top Lp(a)-C interaction variants are all individually associated with Lp(a)-C (**Supplementary Table 12**). Lastly, conditional analysis has been suggested to ensure that the interaction model is not mitigated by a separate third variant that explains the interaction. Thus, we performed conditional analysis on the top 3 interaction models, conditioning on the previously identified variants from single variant analysis (reported in **Supplementary Table 9**) found to be conditionally independently associated with Lp(a)-C in each cohort. As seen in **Supplementary Table 13**, conditional analysis does not fully mitigate any of the identified interaction associations.

### Rare variant association analyses (RVAS)

#### Coding and non-coding grouping schemes

Please refer to the **Supplementary Text** for details on the coding and non-coding grouping schemes used.

#### Statistical Analysis

We tested the association of the aggregate of the aforementioned groupings with each lipid trait using the mixed-model Sequence Kernal Association Test (SKAT) implementation in EPACTS to account for bidirectional effects.^60^ Analyses were adjusted for age, sex, fasting > 10hr, sequencing batch (just used in Estonia), and empiric kinship. Groups with at least 2 rare variants and combined MAF > 0.001 across all aggregated variants in a given cohort were included in meta-analysis. *P* values were meta-analyzed using Fisher’s method. Statistical significance for each RVAS test was based on the number of groups tested and is provided in the headers of **Supplementary Tables 16-25**.

### Mendelian randomization

#### Genetic Instruments

We developed three genetic instruments per cohort. The first instrument used was a genetic risk score, “GRS,” comprised of variants in a ~4MB window around *LPA* (6:158532140-162664257) with sub-threshold significance (p-value < 1×10^−4^), using variant effect sizes from the KIV2-CN conditioned single variant analysis and performing LD clumping in plink using the following parameters: --clump-kb 500 --clump-p1 0.0001 --clump-p2 1 --clump-r2 0.25. This resulted in 399 variants for Lp(a) GRS in FIN, 235 variants for Lp(a) GRS in JHS, 39 variants for Lp(a)-C GRS in JHS, and 49 variants for Lp(a)-C GRS in EST (**Supplementary Table 26-28**). The second instrument used was a “KIV2-CN” score using the directly genotyped or imputed KIV2-CN. The third instrument used was a combined “GRS+KIV2-CN” score combining scores from (1) and (2). Each of the three scores were inverse rank normalized and adjusted such that 1 unit increase in the score is equal to 1SD increase in Lp(a) (or Lp(a)-C, depending on how the instrument was adjusted). The multiplicative factors used to adjust each score are provided in **Supplementary Table 29**. Note: the genetic instruments for EST were not used in MR but are provided as a reference for European Lp(a)-C genetic instruments to compare with those from the JHS African Americans.

#### Phenotypes

Please refer to **Supplemental Text** for details on incident events and subclinical measures used.

#### Statistical analyses

For incident clinical events, a cox proportional hazards test was performed, finding the association between each incident event and each of the genetic instruments as well as observational Lp(a). For the quantitative subclinical measures, linear regression was performed, finding the association between each inverse-rank normalized phenotype and each of the genetic instruments as well as inverse-rank normalized Lp(a) and Lp(a)-C (where available). Covariates used in all analyses included the first 5 principal components of genetic ancestry, age, sex, if the individual was fasting > 10hr. To find the p-value of difference between GRS and KIV2-CN or the genetic instrument and observational phenotype, the following was used:

> *P*(difference)=2*pnorm(-abs(Z)), where Z=b1–b2/(√((SEb1)^2+(SEb2)^2))

Statistical significance was defined for the 9 FIN incident clinical events and 2 JHS subclinical atherosclerosis traits using a Bonferroni significance threshold was based on the number of phenotypes (*P* = 0.0056 and 0.025, respectively).

## REFERENCES

1. Tsimikas, S. & Hall, J.L. Lipoprotein(a) as a potential causal genetic risk factor of cardiovascular disease: a rationale for increased efforts to understand its pathophysiology and develop targeted therapies. J Am Coll Cardiol 60, 716–21 (2012).

2. Utermann, G. The mysteries of lipoprotein(a). Science 246, 904–10 (1989).

3. Berglund, L. & Ramakrishnan, R. Lipoprotein(a): an elusive cardiovascular risk factor. Arterioscler Thromb Vasc Biol 24, 2219–26 (2004).

4. Kraft, H.G., Kochl, S., Menzel, H.J., Sandholzer, C. & Utermann, G. The apolipoprotein (a) gene: a transcribed hypervariable locus controlling plasma lipoprotein (a) concentration. Hum Genet 90, 220–30 (1992).

5. Lanktree, M.B., Anand, S.S., Yusuf, S., Hegele, R.A. & Investigators, S. Comprehensive analysis of genomic variation in the LPA locus and its relationship to plasma lipoprotein(a) in South Asians, Chinese, and European Caucasians. Circ Cardiovasc Genet 3, 39–46 (2010).

6. Lamon-Fava, S. et al. The NHLBI Twin Study: heritability of apolipoprotein A-I, B, and low density lipoprotein subclasses and concordance for lipoprotein(a). Atherosclerosis 91, 97–106 (1991).

7. Austin, M.A. et al. Lipoprotein(a) in women twins: heritability and relationship to apolipoprotein(a) phenotypes. Am J Hum Genet 51, 829–40 (1992).

8. Schmidt, K., Kraft, H.G., Parson, W. & Utermann, G. Genetics of the Lp(a)/apo(a) system in an autochthonous Black African population from the Gabon. Eur J Hum Genet 14, 190–201 (2006).

9. Scholz, M. et al. Genetic control of lipoprotein(a) concentrations is different in Africans and Caucasians. Eur J Hum Genet 7, 169–78 (1999).

10. Mooser, V. et al. The Apo(a) gene is the major determinant of variation in plasma Lp(a) levels in African Americans. Am J Hum Genet 61, 402–17 (1997).

11. Mack, S. et al. A genome-wide association meta-analysis on lipoprotein(a) concentrations adjusted for apolipoprotein(a) isoforms. J Lipid Res (2017).

12. Kraft, H.G. et al. Apolipoprotein(a) kringle IV repeat number predicts risk for coronary heart disease. Arterioscler Thromb Vasc Biol 16, 713–9 (1996).

13. Sandholzer, C. et al. Apo(a) isoforms predict risk for coronary heart disease. A study in six populations. Arterioscler Thromb 12, 1214–26 (1992).

14. Saleheen, D. et al. Apolipoprotein(a) isoform size, lipoprotein(a) concentration, and coronary artery disease: a mendelian randomisation analysis. Lancet Diabetes Endocrinol (2017).

15. Clarke, R. et al. Genetic variants associated with Lp(a) lipoprotein level and coronary disease. N Engl J Med 361, 2518–28 (2009).

16. Kraft, H.G. et al. Frequency distributions of apolipoprotein(a) kringle IV repeat alleles and their effects on lipoprotein(a) levels in Caucasian, Asian, and African populations: the distribution of null alleles is non-random. Eur J Hum Genet 4, 74–87 (1996).

17. Lanktree, M.B. et al. Determination of lipoprotein(a) kringle repeat number from genomic DNA: copy number variation genotyping using qPCR. J Lipid Res 50, 768–72 (2009).

18. Kulkarni, K.R., Garber, D.W., Marcovina, S.M. & Segrest, J.P. Quantification of cholesterol in all lipoprotein classes by the VAP-II method. J Lipid Res 35, 159–68 (1994).

19. Kulkarni, K.R. Cholesterol profile measurement by vertical auto profile method. Clin Lab Med 26, 787–802 (2006).

20. Waldeyer, C. et al. Lipoprotein(a) and the risk of cardiovascular disease in the European population: results from the BiomarCaRE consortium. Eur Heart J 38, 2490–2498 (2017).

21. Handsaker, R.E., Korn, J.M., Nemesh, J. & McCarroll, S.A. Discovery and genotyping of genome structural polymorphism by sequencing on a population scale. Nat Genet 43, 269–76 (2011).

22. Noureen, A., Fresser, F., Utermann, G. & Schmidt, K. Sequence variation within the KIV-2 copy number polymorphism of the human LPA gene in African, Asian, and European populations. PLoS One 10, e0121582 (2015).

23. Handsaker, R.E. et al. Large multiallelic copy number variations in humans. Nat Genet 47, 296–303 (2015).

24. Lim, E.T. et al. Distribution and medical impact of loss-of-function variants in the Finnish founder population. PLoS Genet 10, e1004494 (2014).

25. Kyriakou, T. et al. A common LPA null allele associates with lower lipoprotein(a) levels and coronary artery disease risk. Arterioscler Thromb Vasc Biol 34, 2095–9 (2014).

26. Musunuru, K. et al. From noncoding variant to phenotype via SORT1 at the 1p13 cholesterol locus. Nature 466, 714–9 (2010).

27. Willer, C.J. et al. Discovery and refinement of loci associated with lipid levels. Nat Genet 45, 1274–1283 (2013).

28. Yeang, C., Clopton, P.C. & Tsimikas, S. Lipoprotein(a)-cholesterol levels estimated by vertical auto profile correlate poorly with Lp(a) mass in hyperlipidemic subjects: Implications for clinical practice interpretation of Lp(a)-mediated risk. J Clin Lipidol 10, 1389–1396 (2016).

29. Surakka, I. et al. The impact of low-frequency and rare variants on lipid levels. Nat Genet 47, 589–97 (2015).

30. Li, J. et al. Genome- and exome-wide association study of serum lipoprotein (a) in the Jackson Heart Study. J Hum Genet 60, 755–61 (2015).

31. Lu, W. et al. Evidence for several independent genetic variants affecting lipoprotein (a) cholesterol levels. Hum Mol Genet 24, 2390–400 (2015).

32. Fish, A.E., Capra, J.A. & Bush, W.S. Are Interactions between cis-Regulatory Variants Evidence for Biological Epistasis or Statistical Artifacts? Am J Hum Genet 99, 817–830 (2016).

33. Wood, A.R. et al. Another explanation for apparent epistasis. Nature 514, E3–5 (2014).

34. Roadmap Epigenomics, C. et al. Integrative analysis of 111 reference human epigenomes. Nature 518, 317–30 (2015).

35. Kim, S., Jhong, J.H., Lee, J. & Koo, J.Y. Meta-analytic support vector machine for integrating multiple omics data. BioData Min 10, 2 (2017).

36. Morrison, A.C. et al. Practical Approaches for Whole-Genome Sequence Analysis of Heart- and Blood-Related Traits. Am J Hum Genet 100, 205–215 (2017).

37. Lesurf, R. et al. ORegAnno 3.0: a community-driven resource for curated regulatory annotation. Nucleic Acids Res 44, D126–32 (2016).

38. Liu, Y., Sarkar, A., Kheradpour, P., Ernst, J. & Kellis, M. Evidence of reduced recombination rate in human regulatory domains. Genome Biol 18, 193 (2017).

39. Emdin, C.A. et al. Phenotypic Characterization of Genetically Lowered Human Lipoprotein(a) Levels. J Am Coll Cardiol 68, 2761–2772 (2016).

40. Kettunen, J. et al. Genome-wide study for circulating metabolites identifies 62 loci and reveals novel systemic effects of LPA. Nat Commun 7, 11122 (2016).

41. Guerra, R. et al. Lipoprotein(a) and apolipoprotein(a) isoforms: no association with coronary artery calcification in the Dallas Heart Study. Circulation 111, 1471–9 (2005).

42. Marcovina, S.M., Hobbs, H.H. & Albers, J.J. Relation between number of apolipoprotein(a) kringle 4 repeats and mobility of isoforms in agarose gel: basis for a standardized isoform nomenclature. Clin Chem 42, 436–9 (1996).

43. Jun, G., Wing, M.K., Abecasis, G.R. & Kang, H.M. An efficient and scalable analysis framework for variant extraction and refinement from population-scale DNA sequence data. Genome Res 25, 918–25 (2015).

44. Jun, G. et al. Detecting and estimating contamination of human DNA samples in sequencing and array-based genotype data. Am J Hum Genet 91, 839–48 (2012).

45. Tan, A., Abecasis, G.R. & Kang, H.M. Unified representation of genetic variants. Bioinformatics 31, 2202–4 (2015).

46. Li, B. & Leal, S.M. Methods for detecting associations with rare variants for common diseases: application to analysis of sequence data. Am J Hum Genet 83, 311–21 (2008).

47. Li, H. & Durbin, R. Fast and accurate short read alignment with Burrows-Wheeler transform. Bioinformatics 25, 1754–60 (2009).

48. Van der Auwera, G.A. et al. From FastQ data to high confidence variant calls: the Genome Analysis Toolkit best practices pipeline. Curr Protoc Bioinformatics 11, 11 10 1–11 10 33 (2013).

49. Ganna, A. et al. Ultra-rare disruptive and damaging mutations influence educational attainment in the general population. Nat Neurosci 19, 1563–1565 (2016).

50. Li, H. Toward better understanding of artifacts in variant calling from high-coverage samples. Bioinformatics 30, 2843–51 (2014).

51. Vartiainen, E. et al. Thirty-five-year trends in cardiovascular risk factors in Finland. Int J Epidemiol 39, 504–18 (2010).

52. Howie, B.N., Donnelly, P. & Marchini, J. A flexible and accurate genotype imputation method for the next generation of genome-wide association studies. PLoS Genet 5, e1000529 (2009).

53. Goldstein, J.I. et al. zCall: a rare variant caller for array-based genotyping: genetics and population analysis. Bioinformatics 28, 2543–5 (2012).

54. Delaneau, O., Zagury, J.F. & Marchini, J. Improved whole-chromosome phasing for disease and population genetic studies. Nat Methods 10, 5–6 (2013).

55. Peloso, G.M. et al. Association of low-frequency and rare coding-sequence variants with blood lipids and coronary heart disease in 56,000 whites and blacks. Am J Hum Genet 94, 223–32 (2014).

56. Hoggart, C.J. et al. Control of confounding of genetic associations in stratified populations. Am J Hum Genet 72, 1492–1504 (2003).

57. Libiger, O. & Schork, N.J. A Method for Inferring an Individual's Genetic Ancestry and Degree of Admixture Associated with Six Major Continental Populations. Front Genet 3, 322 (2012).

58. Price, A.L. et al. Principal components analysis corrects for stratification in genome-wide association studies. Nat Genet 38, 904–9 (2006).

59. McLaren, W. et al. Deriving the consequences of genomic variants with the Ensembl API and SNP Effect Predictor. Bioinformatics 26, 2069–70 (2010).

60. Kang, H.M. et al. Variance component model to account for sample structure in genome-wide association studies. Nat Genet 42, 348–54 (2010).

61. Willer, C.J., Li, Y. & Abecasis, G.R. METAL: fast and efficient meta-analysis of genomewide association scans. Bioinformatics 26, 2190–1 (2010).

62. Loh, P.R. et al. Contrasting genetic architectures of schizophrenia and other complex diseases using fast variance-components analysis. Nat Genet 47, 1385–92 (2015).

63. Purcell, S. et al. PLINK: a tool set for whole-genome association and population-based linkage analyses. Am J Hum Genet 81, 559–75 (2007).

